# BioMADE: Predicting Torsades de Pointes from molecular structures through biologically informed representations

**DOI:** 10.64898/2026.05.06.723121

**Authors:** Jose Miguel Acitores Cortina, Martijn C. Schut, Nicholas P. Tatonetti

**Affiliations:** Department of Computational Biomedicine, Cedars-Sinai, West Hollywood, California, USA; Cedars-Sinai Cancer Center, Cedars-Sinai, Los Angeles, California, USA; Translational AI in Laboratory Medicine, Amsterdam UMC, Vrije Universiteit, Amsterdam, The Netherlands

**Author notes:** Contributing authors.

**Keywords:** Torsades de Pointes, Arrhythmias, ADE, ML

## Abstract

Drug-induced arrhythmias, particularly Torsades de Pointes (TdP), pose a significant risk to patient safety and can sometimes have life-threatening outcomes. They remain a major concern in drug development and regulation. Machine learning (ML) has become a powerful tool for analyzing complex biological and chemical datasets, enabling researchers to identify subtle patterns that differentiate safe compounds from those likely to cause dangerous cardiac effects. However, most existing in silico approaches do not sufficiently incorporate biological elements, relying heavily on chemical and structural properties or on computationally expensive simulations.

Here, we introduce BioMADE, a novel ML framework that harnesses small-molecule–protein activity profiles from publicly available datasets to predict TdP risk without requiring exhaustive mechanistic annotation. Activity data from ChEMBL were used to train individual models for each gene, which predict activity values for any given compound. A curated set of arrhythmia-relevant genes was then used to construct a latent biological embedding (BioMADE embedding) for each molecule. We validated the performance of these features in distinguishing biological elements such as ATC3 class, showing superior classification performance compared with representations such as Molformer (lacks biological information) and MACCS (limited chemical properties) (0.85 AUROC vs 0.81 and 0.73, respectively). BioMADE representations served as input to a support vector machine classifier to discriminate TdP-inducing drugs from safe compounds.

BioMADE achieved an AUROC of 0.89 in internal validation, indicating strong predictive performance. Against state-of-the-art models such as ADMEThyst, BioMADE achieved an AUROC of 0.74 on ADMEThyst’s validation set (vs. 0.72 for ADMEThyst). When we combined both approaches, the AUROC reached 0.77.

These results demonstrate that BioMADE provides a scalable, biology-informed, and generalizable approach for predicting drug-induced toxicities. By integrating protein activity profiles into toxicology modeling, our framework highlights the critical role of human biology in adverse drug reaction prediction, an aspect often overshadowed by purely chemical or structural descriptors.

## 1 Introduction

Torsades de Pointes (TdP), a dangerous ventricular tachycardia that can lead to syncope, seizures, and sudden death[1, 2], exemplifies one of the most critical adverse drug reactions in clinical practice. This life-threatening arrhythmia is often caused by drug-induced long QT syndrome (LQTS) and in most cases, preventable. As a well-studied event, it is an ideal target for computational prediction methods. Despite being a known toxicity, many TdP-inducing drugs remain in clinical use, underscoring the urgent need for better predictive tools.

Adverse drug reactions continue to pose significant risks throughout drug development. Drug toxicities represent close to 30% of clinical trial failures, including 24% in phases II and III[3], and many more are withdrawn from the market after approval for unexpected toxicities. Accurately predicting which molecular structures will induce TdP is essential for improving patient safety and reducing costly late-stage failures.

Several approaches have attempted to predict drug toxicities using various methods. Tools like the Therapeutics Data Commons[4] provide frameworks for exploring ADMET properties, with models like ADMETBoost[5] and ADMET-AI[6] demonstrating robust performance, though they lack specificity for individual toxicities. More targeted efforts have addressed cardiac toxicities using the DICTrank dataset[7], including ADMEThyst[8], achieving an AUROC of 0.72 and AUPRC of 0.75 (baseline 0.50), and the DICTrank predictor by Seal et al.[9], which used Mordred[10] properties and obtained an AUPRC of 0.93 (baseline 0.72). Both approaches have limitations: their representations remain mostly chemical, overlooking substantial aspects of human biology, or they address broad cardiotoxicity rather than TdP specifically.

Molecular structures contain extensive information about compounds, including physical properties like polarity and solubility, as well as chemical properties and biological activity[11]. Due to the recent advent of foundation models and their ability to generalize and learn from large amounts of data, toxicity research has incorporated them to extract molecular information by training them on billions of small molecules (e.g., MolFormer[12], UniMol2[13]). These models can generate more complete molecular representations than their predecessors, such as MACCS or Morgan fingerprints. These representations have proven useful for downstream property prediction tasks.

However, to achieve accurate predictions, it is necessary to effectively bridge molecular chemistry with human biology, as translating these representations into predictions of biological toxicity remains difficult. The largest dataset of small molecules, the chemical universe database GDB-17, contains 166.4 billion possible small molecules [14], yet fewer than 220 million substances have been registered by the Chemical Abstracts Service [15], and only approximately 2.4 million have experimentally confirmed bioactivity [16]. Of these, around 21,000 have entered clinical trials [17], and the FDA has approved roughly 1,300 unique active ingredients [18]. Given the rarity of bioactivity, the amount of relevant medical information that can be extracted from representations generated by generalist foundation models is limited. There is a critical need to guide these powerful models to represent biologically relevant chemical space, specifically, the molecular features governing drug-target interactions in human cardiac physiology.

There are a few TdP datasets and approaches for predicting TdP or QT prolongation. Datasets include the drug-induced QT prolongation atlas (DIQTA[19]), and Crediblemeds[20], which categorizes drugs by known, possible, and conditional TdP risk. Molformer-XL-CNN[12] used DIQTA, achieving an AUROC of 0.82. Yap et al. [21] used the Crediblemeds dataset from 2004 and achieved a precision of 0.97 for TdP-causing agents and 0.84 for non-TdP-causing agents; however, the dataset does not reflect current classifications, and applying their approach to current data would likely yield lower performance. Other methods for predicting TdP include experimental data and complex software simulations[22–24].

In this work, we introduce Biologically Informed Chemical Models for ADE prediction (BioMADE), a framework that predicts toxicities from molecular structures and captures relevant biology. This framework consists of two steps: first, we train a set of gene-drug interaction models to generate a bioactivity representation for each drug, where each gene is treated as a feature. The second step uses these relevant embedding features to predict TdP. We achieved an AUROC of 0.91 (± 0.05) in our internal validation. Additionally, BioMADE outperforms SOTA models like ADMEThyst on DICTrank (0.72(± 0.07) vs 0.77(± 0.06) AUROC).

## 2 Results

### 2.1 Data

Our data pipeline integrated three primary sources to comprehensively capture drug-induced long QT syndrome and Torsades de Pointes. The first source was the Credible Meds Torsades de Pointes dataset[20], which provides clinically validated information on drugs associated with prolongation of the QT interval and risk of ventricular arrhythmias. The second source was the DICTrank dataset[7], which contains a large set of cardiotoxic drugs and controls; we used the provided controls. Lastly, we used ChEMBL35[16], a large-scale bioactivity database containing curated compound, target, and bioactivity data from the scientific literature. By combining these complementary datasets, we established a comprehensive framework for identifying and characterizing potentially cardiotoxic compounds related to QT prolongation and life-threatening arrhythmias.

### 2.2 Approximation of gene-drug activities

We selected genes with more than 100 reported interactions in ChEMBL and trained a Chemprop model for each to approximate gene–drug interaction activities as a pChEMBL value. Chemprop is a machine learning library for chemical property prediction; its models consist of four modules: a local feature encoding function, a directed message passing neural network (D-MPNN) to learn atomic embeddings from local features, an aggregation function to combine atomic embeddings into molecular embeddings, and a standard feedforward neural network (FFN) for transforming molecular embeddings into target properties [25]. We leveraged this architecture to predict the interaction between a small molecule and a target protein using pchembl values as the metric. By creating one model per available gene, we eliminate the need to encode the protein to calculate bioactivity values.

As shown in Figure 1, our models achieved a wide range of RMSEs and MAEs, which were correlated with the number of activities reported in ChEMBL. Of these, 1,274 models achieved an RMSE below 1 (corresponding to predictions within one order of magnitude of the true value), and 145 achieved an RMSE below 0.5. A notable example is KCNH2, which plays a critical role in cardiac repolarization; it has nearly 10,000 activities recorded in ChEMBL35, and its model achieved an RMSE of 0.55.

**Fig. 1.**
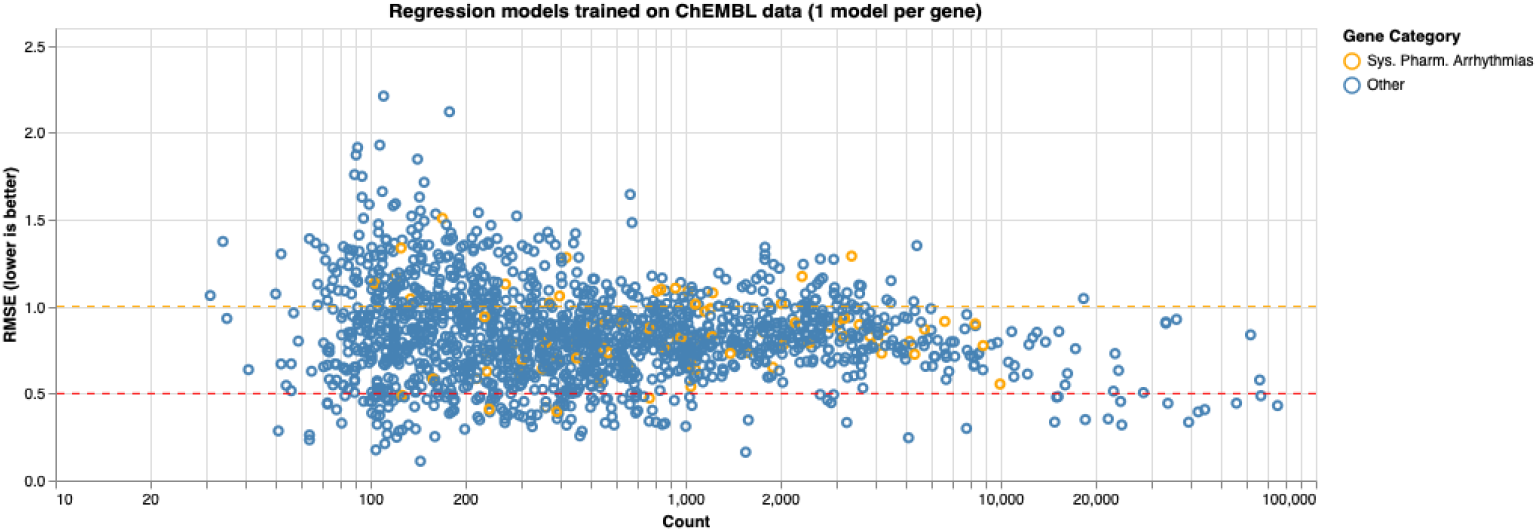
shows a scatter plot of the RMSEs of all the Chemprop models trained for each gene with at least 100 interactions in ChEMBL.

Other genes previously associated with long QT syndrome, such as EGFR, HTR2A, OPRS1, and MAPK14, showed RMSEs below 1, each with more than 5,000 recorded activities.

Each of these predictions represents a feature in the BioMADE molecular representation. To validate the hypothesis that BioMADE embeddings capture biological information better than more general representations, we compared the performance of MACCS fingerprints, Molformer embeddings, and BioMADE embeddings to predict whether two molecules share the same ATC3 classification. We used all available drugs in DrugBank and generated all possible pairs, assigning a label of zero if they do not share an ATC3 class and one if they do. We then trained a logistic regression model to evaluate each pair. As shown in Figure 2, BioMADE outperformed both alternative representations, achieving an AUROC of 0.86.

**Fig. 2.**
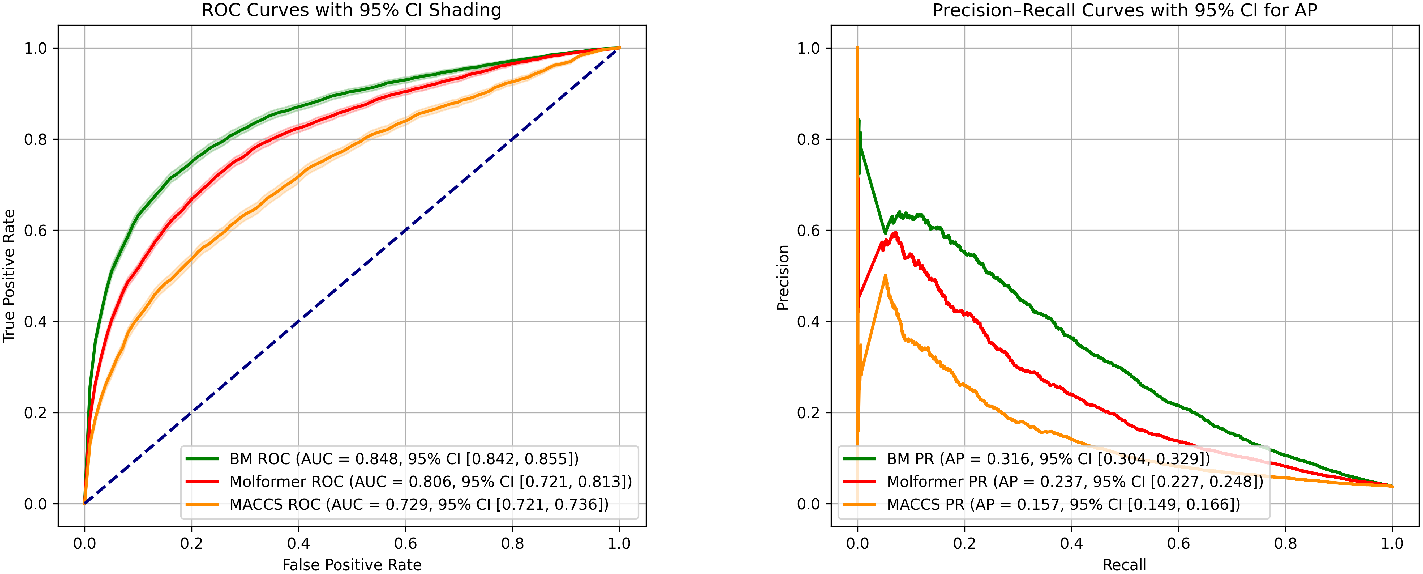
shows the ROC and PR curves resulting from the ATC3 classification of drug pairs by a logistic regression trained on BioMADE embeddings, Molformer embeddings and MACCS representations.

To generate a molecular representation, we initiated a feature selection process to identify the optimal gene set. We focused on biological pathways, reasoning that selecting genes involved in TdP-relevant functions should yield better predictive performance. We benchmarked all pathways available in Reactome for the task of predicting TdP, as well as the genes implicated in the mechanism of arrhythmias as evaluated by Berger et al. in their work on the systems pharmacology of arrhythmias [26]. We split the dataset into training and test sets, ensuring that no two drugs sharing the same ATC4 category appeared in different sets to minimize data leakage. Results are shown in Figure 3.

**Fig. 3.**
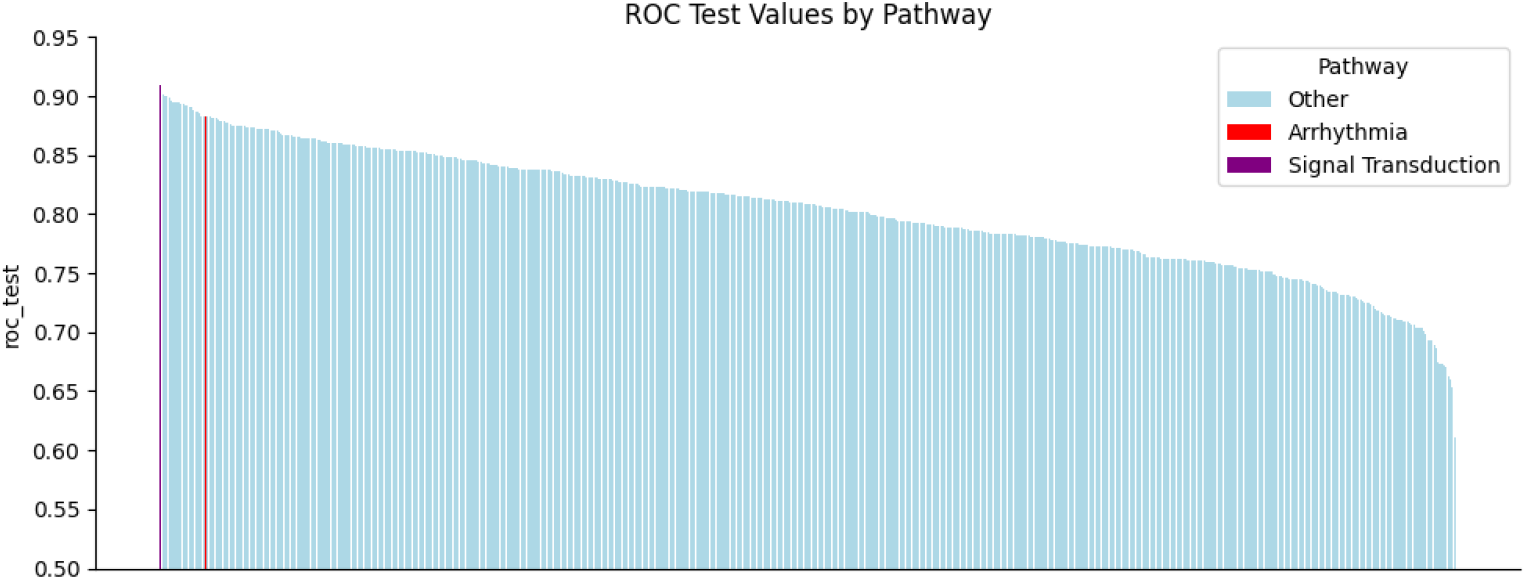
shows the AUROC of all gene groups, grouped by pathways extracted from Reactome and the additional groups identified. Signal Transduction and Systems Pharmacology of Arrhythmias are highlighted at the top.

**Fig. 4.**
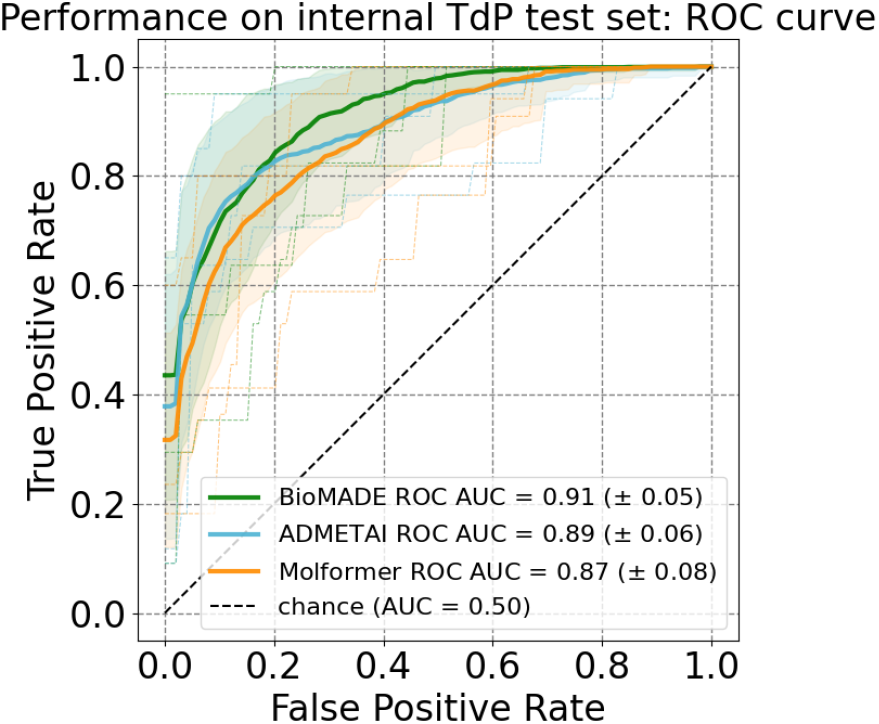
shows the comparison of performance between two state-of-the-art models and BioMADE. Molformer embeddings with an XGBoost classifier achieved an AUROC of 0.87 (± 0.08). ADMET-AI with an XGBoost classifier head achieved 0.89 (± 0.06) AUROC. BioMADE, using Signal Transduction pathway embeddings and a logistic regression head, achieved 0.91 (± 0.05) AUROC.

**Fig. 5.**
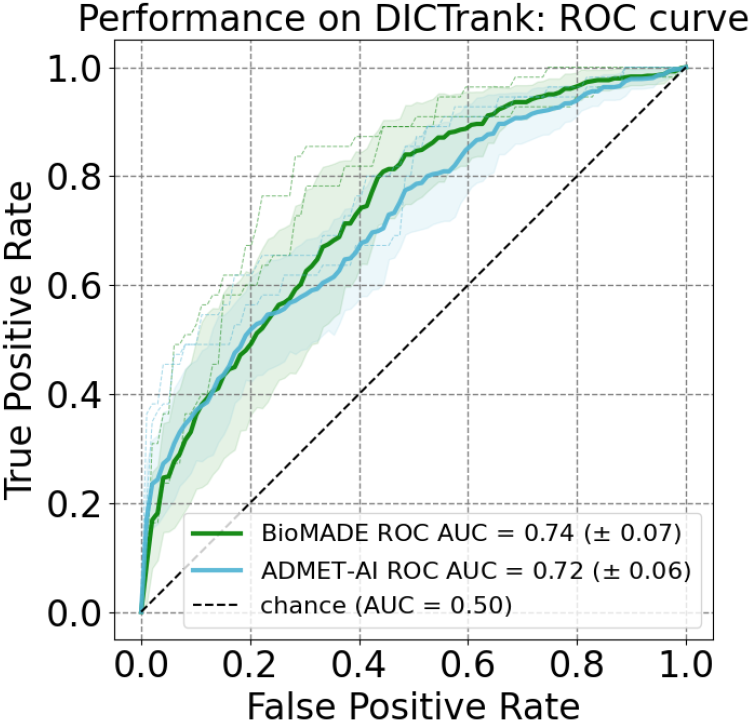
shows the receiver operating characteristic curve for external validation on DICTrank. ADMET-AI with an XGBoost classifier head achieved 0.72 (± 0.06) AUROC, and BioMADE using the Systems Pharmacology of Arrhythmias gene set achieved 0.74 (± 0.07) AUROC.

### 2.3 Predicting TdP from estimated activities

Our dataset for predicting TdP comprised all drugs with known or possible TdP risk according to the CredibleMeds [20] dataset as positive cases, and a set of controls from the DICTrank [7] dataset with no known risk of cardiotoxicity. We selected two main gene groups to predict TdP risk. The first group consisted of genes related to the systems pharmacology of arrhythmias, as identified by Berger et al. [26]. Training a support vector machine (SVM) using this gene set achieved an AUROC of 0.88. Although only 78 genes overlapped with those for which ChEMBL prediction models were available, this smaller set still produced competitive results, suggesting that arrhythmia-associated genes carry higher predictive value for TdP.

The second gene set comprised genes involved in the Signal Transduction pathway, with 647 overlapping genes and the highest AUROC of 0.91. We chose these two gene sets for our final analyses. The results for all the pathways tested can be found in supplemental materials.

Finally, we tested our model using all available gene representations with RMSE scores of 1 or lower. This approach performed slightly worse than our best pathway-specific model. Detailed results are reported in Table 2.

### 2.4 Internal validation

We then compared BioMADE against two state-of-the-art models for molecular property prediction and cardiotoxicity prediction: Molformer and ADMET-AI. Both models were trained using the same data splits and evaluation protocol as BioMADE to ensure a fair comparison. As shown in Table 1, BioMADE was the best-performing model for predicting TdP on our dataset.

**Table 1.** Results of state-of-the-art models on our internal validation set.

| Model | AUROC | AUPRC |
| --- | --- | --- |
| ADMET-AI | 0.89 ($\pm 0.06$) | 0.82 ($\pm 0.10$) |
| Molformer | 0.87 ($\pm 0.08$) | 0.80 ($\pm 0.09$) |
| BioMADE | 0.91 ($\pm 0.05$) | 0.83 ($\pm 0.10$) |

**Table 2.** Results of our models in the internal validation dataset.

| Gene set | Model | AUROC test | CI95 | AUROC train | AUPRC |
| --- | --- | --- | --- | --- | --- |
| Systems pharmacology of arrhythmias | LR | 0.88 | .011 | 0.91 | 0.76 |
|  | SVM | 0.88 | .011 | 0.92 | 0.67 |
|  | RF | 0.81 | .091 | 1.00 | 0.59 |
|  | XGB | 0.79 | .121 | 1.00 | 0.58 |
| Signal Transduction | LR | 0.91 | .009 | 0.99 | 0.83 |
|  | SVM | 0.91 | .012 | 0.93 | 0.80 |
|  | RF | 0.84 | .085 | 1.00 | 0.69 |
|  | XGB | 0.82 | .121 | 1.00 | 0.69 |
| All available genes | LR | 0.90 | .010 | 0.99 | 0.82 |
|  | SVM | 0.90 | .012 | 0.92 | 0.69 |
|  | RF | 0.86 | .057 | 1.00 | 0.70 |
|  | XGB | 0.80 | .109 | 1.00 | 0.57 |

The baseline AUPRC was 0.3 on average.

### 2.5 External validation

One of the pillars of our research was the model’s generalizability. Therefore, we performed external validation on unseen data using intuitive gene sets and benchmarked our model against state-of-the-art approaches.

#### 2.5.1 DICTrank

We trained and tested BioMADE with the Systems Pharmacology of Arrhythmias gene set in two configurations. First, following the data split of ADMET-AI, BioMADE out-performed ADMEThyst, achieving an AUROC of 0.74 on the validation set (vs. 0.72 for ADMEThyst). Combining both approaches yielded an AUROC of 0.77. Our second DICTrank validation followed the data split of the DICTrank predictor. BioMADE performed comparably with DTpred on their set. Detailed results are reported in Table 3.

**Table 3.** Results of the three models on different splits of the DICTrank dataset.

| Model | split | AUROC | AUPRC |
| --- | --- | --- | --- |
| ADMET-AI | ADMETHyst | 0.72 ($\pm 0.06$) | 0.74 ($\pm 0.08$) |
| BioMADE | ADMETHyst | 0.74 ($\pm 0.07$) | 0.76 ($\pm 0.08$) |
| ADMET-AI + BioMADE | ADMETHyst | 0.77 ($\pm 0.06$) | 0.74 ($\pm 0.08$) |
| DT pred (Mordred) | DTpred | 0.84 | 0.93 |
| BioMADE (XGB)<sup>1</sup> | DTpred | 0.84 | 0.93 |

Baseline AUPRC for the ADMETHyst data split is 0.5. Baseline AUPRC for the DICTrank pred data split is 0.72.
<sup>1</sup>This model was trained on an extended cardiomyopathy gene set from WashU Medicine Pathology Services.

## 3 Discussion

This study introduces BioMADE, a novel machine learning framework that leverages biologically informed molecular representations to predict drug-induced Torsades de Pointes (TdP) risk. Our approach addresses a critical limitation in current computational toxicology methods: the inability to capture biologically relevant molecular features that drive adverse drug reactions. By integrating gene–drug interaction data from ChEMBL with pathway-informed feature selection, BioMADE creates molecular embeddings that encode mechanistic information relevant to cardiac arrhythmogenesis.

BioMADE’s performed better than state-of-the-art approaches, including ADMET-AI and Molformer, demonstrating the value of incorporating human biology into molecular representation learning. Our model achieved an AUROC of 0.91 in internal validation, and 0.74 AUROC on the ADMEThyst split of DICTrank, and 0.84 AUROC on a different DICTrank partition, BioMADE outperformed purely structure-based methods while maintaining interpretability through its biologically grounded feature space.

The success of the ATC3 classification task highlights BioMADE’s ability to capture pharmacologically relevant molecular patterns that extend beyond simple structural similarities.

A notable finding is that the Signal Transduction pathway gene set (647 genes) outperformed the more disease-specific Systems Pharmacology of Arrhythmias set (78 genes), achieving an AUROC of 0.91 versus 0.88. Although the arrhythmia-specific genes are more directly implicated in cardiac electrophysiology, several factors may explain the superior performance of the broader pathway. First, the Signal Transduction gene set provides a substantially richer feature space, which generates a richer feature space. Second, signal transduction pathways encompass upstream regulators of ion channel expression and trafficking—processes such as PKA- and PKC-mediated phosphorylation of hERG channels, MAPK-dependent transcriptional regulation of *K*_*V*_ and *Na*_*V*_ subunits, and PI3K/Akt-mediated modulation of *I*_*Kr*_ current density— that are not captured by a narrow focus on the channels themselves. Third, TdP liability frequently arises from off-target pharmacology; drugs may perturb signaling cascades that indirectly alter cardiac repolarization reserve without directly binding canonical arrhythmia targets. This observation supports the broader thesis of BioMADE that encoding a drug’s activity across a wide panel of biologically relevant targets yields representations that capture the pharmacological complexity underlying adverse drug reactions.

Another important finding from our results points towards the model selection. The ensemble methods exhibited clear signs of overfitting. In contrast, simpler linear classifiers achieved the highest generalization performance (AUROC of 0.91 and 0.88, respectively) without a big train–test discrepancy. This suggests that BioMADE embeddings already encode biologically relevant nonlinear relationships between molecular structure and TdP risk in a form that is approximately linearly separable. Therefore, additional model complexity provides no benefit and instead fits noise in the training data. This finding highlights that the primary value of BioMADE lies in the representation itself rather than in the downstream classifier, and that the embedding can be deployed with lightweight, interpretable models suitable for regulatory and clinical decision-support contexts.

There are several limitations that should be mentioned, our models rely on publicly available data that may be incomplete or biased toward well-studied targets, potentially limiting generalizability to novel chemical compounds. Second, although BioMADE captures gene-level interactions, it does not explicitly model protein structure, protein–protein interactions, or post-translational modifications that may influence drug–target binding. Third, the current framework predicts binary TdP risk and does not account for dose dependence or individual patient variability, both of which are critical in clinical practice. Fourth, our model does not account for drug-drug interactions, which can be the culprit of many adverse drug reactions.

Future work should explore expanding the ADEs studied, incorporating multiomics data (e.g., transcriptomics, proteomics) to enrich the biological embedding, developing tissue-specific models that account for cardiac-specific gene expression patterns, and extending the approach to predict dose-dependent toxicity and drug–drug interactions. Integration with experimental assay data, such as those generated under the CiPA initiative, could further enhance predictive accuracy.

In conclusion, BioMADE represents a significant advance in computational toxicology by demonstrating that biologically informed molecular representations improve the prediction of drug-induced adverse effects while introducing interpretability. Our work highlights the importance of incorporating human biology into computational drug safety assessment, moving beyond purely chemical perspectives to account for the biological complexity that ultimately determines therapeutic outcomes.

## 4 Methods

### 4.1 Training of Chemprop models

We extracted bioactivity data from ChEMBL 35 to construct predictive molecular features via Chemprop-based modeling. Bioactivities were grouped by target gene, and only targets with more than 100 unique compound–activity entries were retained to ensure sufficient training data. For each qualifying gene, a separate Chemprop model was trained using Python. The model architecture consisted of a Chemprop BondMessagePassing layer to extract molecular structure information, followed by a MeanAggregation layer to generate fixed-length molecular embeddings, and a feedforward neural network (RegressionFFN) to predict the pChEMBL value. Targets were normalized prior to training. Training was performed using the Adam optimizer with batch normalization, GPU acceleration, and early stopping with a patience of 5 epochs. After model selection, each model was retrained on 100% of the data to generate the final molecular representations.

### 4.2 Embedding construction and feature selection

For each drug, an embedding vector was generated by passing the molecule through every trained gene-specific model. The output of each model served as a single feature, resulting in a feature vector in which each dimension corresponded to one target gene. A subset of features was selected based on prior knowledge of biological pathways and published studies implicating specific targets in cardiac electrophysiology and arrhythmogenesis, as detailed in the Results section. This biologically informed selection aimed to enhance model interpretability and predictive relevance for TdP risk.

### 4.3 TdP Risk Prediction Models

The selected features were used to train models to predict TdP risk, with the CredibleMeds and DICTrank classifications serving as references. Data were split into training and test sets, stratified by ATC level 3 (ATC3) classification to prevent information leakage between the sets. We evaluated multiple machine learning algorithms:

- Extreme Gradient Boosting (XGBoost)
- Logistic Regression (LR)
- Support Vector Machine (SVM)
- Neural Network (NN)
- Random Forest (RF)

Model performance was assessed using standard metrics, including accuracy, precision, recall, F1-score, and area under the receiver operating characteristic curve (ROC-AUC).

## 5 Data

This study integrates data from four complementary sources to construct biologically informed molecular representations and evaluate their predictive capacity for drug-induced QT-interval prolongation and Torsades de Pointes.

### 5.1 ChEMBL 35 Bioactivity Data

Bioactivity and compound metadata were obtained from ChEMBL version 35 [16], a manually curated database of bioactive molecules with drug-like properties. We extracted all records from the activities table that had an associated pChEMBL value, which groups experimentally measured potency metrics (*IC*_50_, *EC*_50_, *K*_*i*_, *K*_*d*_) for compounds tested against a diverse panel of macromolecular targets, including voltage-gated ion channels (e.g., hERG/KCNH2, *Na*_*V*_ 1.5/SCN5A, *Ca*_*V*_ 1.2/CACNA1C) and G-protein-coupled receptors relevant to cardiac electrophysiology. We kept activities derived from single-protein target assays on human or human-ortholog targets. After filtering, the dataset comprised activities across 1,546 unique gene targets, each with at least 100 compound–activity entries, totaling approximately 2 million bioactivity records.

We downloaded all the activities (from the activities table) with available pchembl value from ChEMBL35.

### 5.2 CredibleMeds QT Interval Risk Classification

The CredibleMeds database [20] provides clinically validated, expert-curated lists of medications associated with QT-interval prolongation and TdP risk, stratified into “Known Risk,” “Possible Risk,” and “Conditional Risk” categories based on the strength of clinical and pharmacovigilance evidence. The QTDrug list was extracted through the CredibleMeds API. Drugs classified as “Known Risk” or “Possible Risk” were selected as positive cases for TdP prediction. Compound identifiers were harmonized to canonical SMILES and InChIKey representations and cross-referenced against ChEMBL and DrugBank entries using name, synonym, and structural matching. The final positive set comprised 69 Known risk drugs and 154 possible risk drugs.

### 5.3 Drug-Induced Cardiotoxicity Rank (DICTrank) Dataset

The DICTrank dataset [7], curated by the U.S. FDA’s National Center for Toxicological Research, provides a quantitative ranking of 1,318 FDA-approved drugs by their likelihood of inducing cardiotoxic effects. Rankings are derived from integrated evidence spanning drug labeling, pharmacovigilance reports, and published literature, with standardized annotations for cardiotoxic endpoints including QT prolongation, arrhythmia, and heart failure. Compounds categorized as “No Concern” for cardiotoxicity were selected as negative controls for the TdP prediction task. DICTrank entries were matched to ChEMBL compounds using InChIKey identifiers, yielding 169 control compounds after excluding any overlap with the CredibleMeds positive set.

### 5.4 Reactome Pathway Annotations

Pathway–gene membership data were obtained from Reactome, a peer-reviewed, open-access knowledge base of biological pathways and processes. All human pathway definitions were downloaded in JSON format and parsed to extract gene sets for each pathway. These annotations were used for biologically informed feature selection, enabling the construction of pathway-specific BioMADE embeddings as described in the Results section. A total of 2751 Reactome pathways were evaluated, spanning signaling, metabolism, immune function, and developmental processes.

## Data availability

Data from ChEMBL is publicly available and can be found at their site (https://chembl.gitbook.io/chembl-interface-documentation/downloads).

Data from DICTrank is publicly available and can be found at the FDA website (https://www.fda.gov/science-research/bioinformatics-tools/drug-induced-cardiotoxicity-rank-dictrank-dataset) Long QT Data from CredibleMeds is available for registered users at their website (https://www.crediblemeds.org/)

## Code availability

The code will be made available after peer reviewed publication.

## Acknowledgements

This work was supported by NIH R35GM131905.

## Author contributions

J.M.A.C. conceptualized the study, developed the methodology, wrote the software, performed the formal analysis, curated the data, created the visualizations, and wrote the original draft. M.C.S. contributed to the methodology and reviewed and edited the manuscript. N.P.T. conceptualized the study, supervised the research, administered the project, and reviewed and edited the manuscript.

